# Using indication embeddings to represent patient health for drug safety studies

**DOI:** 10.1101/737049

**Authors:** Rachel D. Melamed

**Affiliations:** Biomedical Data Science, University of Chicago, 900 E 57 St, Chicago, IL, USA; Biology, University of Massachusetts, Lowell, 198 Riverside St, Lowell, MA, USA

**Keywords:** representation, causal inference, cohort studies, embeddings, dimensionality reduction

## Abstract

**Objective:** The electronic health record is a rising resource for quantifying medical practice, discovering adverse effects of drugs, and studying comparative effectiveness. One of the challenges of applying these methods to health care data is the high dimensionality of the health record. Methods to discover effects of drugs in health data must account for tens of thousands of potentially relevant confounders. Our goal in this work is to reduce the dimensionality of the health data with the aim of accelerating the application of retrospective cohort studies to this data.

**Materials and Methods:** Here, we develop indication embeddings, a way to reduce the dimensionality of health data while capturing information relevant to treatment decisions. We evaluate these embeddings using external data on drug indications. Then, we use the embeddings as a substitute for medical history to match patients, and develop evaluation metrics for these matches.

**Results:** We demonstrate that these embeddings recover therapeutic uses of drugs. We use embeddings as an informative representation of relationships between drugs, between health history events and drug prescriptions, and between patients at a particular time in their health history. We show that using embeddings to match cohorts improves the balance of the cohorts, even in terms of poorly measured risk factors like smoking.

**Discussion and Conclusion:** Unlike other embeddings inspired by word2vec, indication embeddings are specifically designed to capture the medical history leading to prescription of a new drug. For retrospective cohort studies, our low-dimensional representation helps in finding comparator drugs and constructing comparator cohorts.

## Introduction

The Electronic Health Record (EHR) is an increasingly complete record of human health and medical practice. Recent studies have used the EHR to evaluate variation in drug treatment decisions,[1,2] identify symptoms related to diseases,[3,4] and to annotate the medical conditions for which drugs are prescribed, known as the indications for a drug.[5,6] One major area of research repurposes EHR data to uncovers adverse effects among people taking a drug, or perform comparative effectiveness studies, using the cohort study design as a way to complement randomized trials.[7–9]

All of these efforts must confront the high dimensional nature of health data: there are tens of thousands of ICD codes specifying particular diagnoses, and thousands of common drugs, alongside other medical events such as procedures and tests. The most straightforward way to code such discrete data for use in an algorithm results in a high-dimensional vector that is sparse, such as a one-hot (dummy variable) vector for each ICD code or drug (Figure 1A). Furthermore, these vectors lack information: the sparse, high-dimensional vector that represents a diagnosis for diabetes will be no nearer to the vector for insulin, than to the vector for acne. To address this issue, a number of studies have explored alternative lower-dimensional representations of health data.

**Figure 1.**
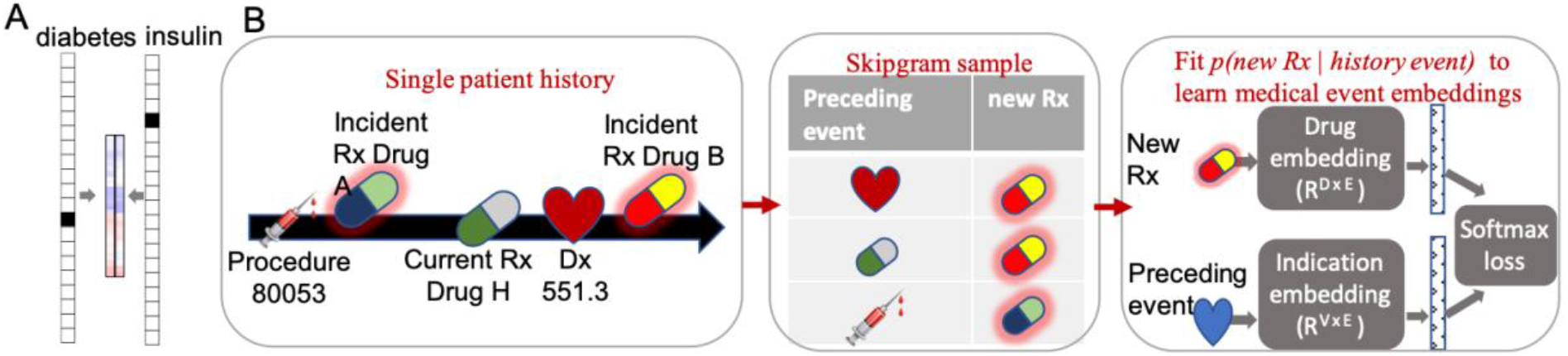
A. Illustration of transformation of embeddings. Coded data can be computed on by representing each code as a one-hot vector with tens of thousands of dimensions. Embeddings allow us to represent the code as a small dense vector of under 100 dimensions. B. Outline of skipgram creation and simplified neural network to create indication embeddings. A patient history is parsed into skip-grams, which are pairs of a new drug, and a medical code that preceded that drug. Each skip-gram is fed into the neural network, which over many skip-gram examples, is trained on an auxiliary task of using the embedding vectors to predict which drug will follow an observed medical code.

In particular, Miotto[10] proposed autoencoders to create dense vector summaries of patient health that predicted disease incidence. Choi[11] adapted the Word2Vec method to medical histories. This method, originally popularized for representing natural language[12] transforms the high-dimensional sparse representation of a word to a low-dimensional vector such that words that are used together have closer vectors. These representations have shown success in the task of phenotyping patient cohorts.[13,14] Thus, embeddings have become a popular way to represent disease, but so far, have not been used to contribute to the cohort study literature.

In this work, we describe a new strategy to create embeddings that represent the relationship between medical history and the first prescription of a drug. This medical history could include diseases that are indications for prescription of a drug, or drugs that act to ameliorate the side effects of other drugs (such as potassium supplements for diuretic prescriptions), or procedures that require prescription of a drug (such as colonoscopy preparations). By creating these embeddings, we aim to capture not just patient state, but the variation in patient state that leads to drug prescription.

We then assess how well our embeddings recapitulate de facto drug indications. While databases such as UMLS[15] and MEDI[16] codify well-established uses for drugs, these may not reflect off-label uses and changes in medical practice. Our method provides a data-driven way to identify such prescription choices. But rather than simply classifying a approved indications, we identify a richer context for medication use.

Finally, we explore the application to the important task of conducting drug safety studies using observational data. Comparative cohort studies are a major tool for assessing drug safety, but these studies require appropriate adjustment for confounding variables. Propensity score methods adjust for confounders by estimating the effect of these confounders on treatment assignment. Confronting the high dimensionality of the health record, with tens of thousands of potential confounders, recent studies have proposed improvements to the propensity score, including large-scale regularized estimation, and automatic determination of relevant confounders.[8,17–19]. Failure to create comparable cohorts can add bias and noise to effect estimates.[20,21] Coarsened Exact Matching (CEM) [22] is one way to create more comparable cohorts, but it requires experts to select a few important variables for matching, and it is not well suited to the context of sparse, high-dimensional data.

We examine the utility of our embeddings for constructing comparator cohorts. The first step in constructing such cohorts typically involves picking a comparable drug to the drug of interest. We assess the use of drug embeddings to identify the most comparable drugs: that is, the drugs given to patients with the most similar health backgrounds. The second step in constructing cohorts is to select a control set of people taking the comparator drug who are roughly similar to the treated group. Since the embeddings represent medical events leading up to drug prescription, we explore the use of embeddings to summarize the health status of each patient. Patients with the most similar health status embeddings should not only have similar likelihood of being prescribed the drug of interest, but they may also be prescribed that drug for the same reason.

Our embeddings create a representation of medical history centered on provider treatment choices. We expect that our approach will enable researchers to more rapidly conduct cohort studies and discover characteristics of medical practice.

## Methods

All methods described below were applied to the Marketscan IBM claims data set, which contains NDC-coded prescriptions, ICD-9 coded diagnoses, and CPT/HCPCS procedures, each time-stamped. We match NDC codes to drug generic names using the MarketScan RED BOOK™ Supplement (includes variables related to drug prescription). Both inpatient and outpatient data were combined. Because of the switch from ICD-9 to ICD-10 in 2015, data from before 2015 was used for most of the analysis, but separately, ICD-10 embeddings were created using data from October 2015 onward. Code to create most figures, tables, and statistics is available at https://github.com/RDMelamed/indication-embeddings

### Generation of indication embeddings

To create the indication embeddings, we adapt the method discussed in Mikolov.[12] In that approach, each word in a sentence can be a *label*, and a randomly chosen *context* window size (number of words before and after) around the label word is chosen. Each word in the window forms a *context* for the *label.* Skip-grams, which are pairs of *(context, label)* are the input to the learning procedure, which then creates embeddings that maximize the likelihood of this data. For our application, we create a number of changes to this procedure (Figure 1B). Only new drug prescriptions are *labels*, and events (ICD-9 codes, drug prescriptions, or procedures) preceding a new drug prescription are *context* for the new drug. Therefore, unlike in Word2Vec, there is an asymmetry between context and label. In another difference, medical data, unlike text data, has the element of time: we are more interested in events that happen soon before a prescription. Mikolov. et al., implemented this idea by selecting words in randomly chosen windows up to 10 words around the training word. To adapt this idea to the medical setting, our context window is based on selecting a random number of weeks before prescription, by drawing from half-normal distribution with a standard deviation of 40 weeks. As well, some patients have more data than other patients, which can result in very sick and densely observed patients dominating the distribution of skip-grams. So, when patients have multiple events in two-month period, we randomly select one of these events rather than creating a skip-gram pair for each event. As in many word2vec implementations, this sampling is weighted to downsample the most frequent codes. To explore possible effects of design decisions such as the size of the context window, the down-sampling of common codes, and the choice of a two-month window, we create embeddings for a variety of settings of these parameters, showing no major effect on the results described below (Supplementary Table 1). In addition to the ICD-9 embeddings, we create a separate embedding for the portion of our data from October 2015 through the end of 2016, that includes ICD-10 codes. The ICD-10 embedding is not used in the primary results of this paper, as most of our data is encoded in ICD-9, but this embedding is made available as a resource, as ICD-10 is the new standard.

Given the set of skip-grams generated above, we train a simple neural network to create embeddings that best predict the new-drug labels, using the standard word2vec setup (Figure 1B). For the pair of history event *h* (procedure, drug, or diagnosis) and new prescription, *d*, the current indication embedding for the history event, Θ_*h*_, is retrieved, and passes through a dropout layer to create a noisy version of the embedding, 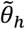. Then, we use a softmax output layer, that has a vector of parameters Φ, for each possible new prescription that can follow a history event. We use Keras with TensorFlow to maximize the softmax function 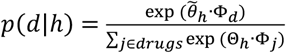 with respect to the parameters Θ and exp Φ. As usual, this proceeds by maximizing the negative log of this function, with a regularization penalty (L2 regularization). We use the usual method of cross-validated hyperparameter selection to select embedding size, L2 regularization parameter, dropout rate, and learning rate, resulting in creating 50-dimensional embedding vectors. The vectors Θ comprise the indication embeddings, which are finally normalized by dividing by the L2 norm so that all embeddings have a norm of 1.

### Visualizations of embeddings

For Figures 2A and 6 the UMAP package from https://umap-learn.readthedocs.io/ is used to map each 50-dimensional embedding vector to a 2-dimensional space. For Figure 2B and 4, the embedding vectors for each code in a CCS category, disease category, or drug category are averaged together, after filtering out codes that appeared in fewer than 20,000 individual patients. For Supplementary Figure 1 the disease embeddings closest to any antidepressant are clustered hierarchically. Because many disease groups are similar (ie, ADHD and Learning Disorder), Figure 4 is a simplified version of Supplementary Figure 1 where the diseases are clustered using K-means clustering (K = 10) and the single disease nearest the centroid of each cluster is chosen to represent that cluster of diseases (ie, Tics Tourettes standing in for Conduct Disorder, Autism, Learning Disorder, Unspecified Childhood Psychoses, ADHD).

**Figure 2.**
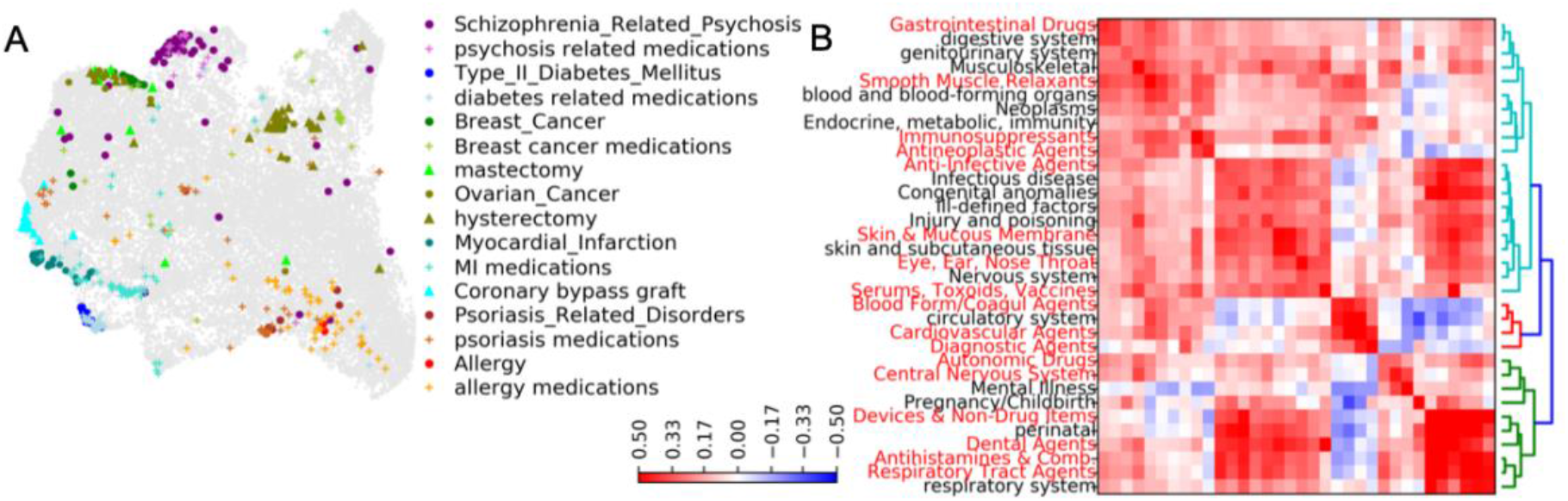
Indication embedding overview. A. Each point is one event (prescription, diagnosis code, or procedure code), visualized with UMAP. Selected related sets of events are highlighted. Circles are ICD-9 codes, + symbols are medications, and triangles are procedures. B. We create groupings of disease codes (black labels) and drug codes (red labels). For each pair of such groupings, we measure the average distance between codes, creating a symmetric distance matrix (color scale).

### Evaluation of embedding vectors

We use the dot product (cosine distance) between a drug’s embedding and an ICD-9 code’s embedding vector as a measure of distance: close drugs and diagnoses imply similar health context. To evaluate the quality of our embedding vectors, we ask whether the closest drugs to an ICD-9 code are the therapies associated with that ICD-9 code (and vice versa), as recorded in MEDI.[16] We use MEDI as our gold-standard for true therapeutic uses. The 10,912 reported therapeutic relationships in the MEDI High Precision Set that overlap with the drugs and ICD-9 codes in our data form our gold-standard positive examples, and 752,812 relationships between the same drugs and ICD-9 codes, where the relationship does not appear in any MEDI database, comprise our negative examples. Then, we can create an ROC curve for the performance of embedding vector distance for predicting indication relationships. We compare performance against a baseline method: co-occurrence of drug with the ICD-9 code preceding the drug. Our baseline co-occurrence method uses two-by-two contingency tables for the co-occurrence of drug and ICD-9 codes. Comparing the observed number of co-occurrences to the number expected given the marginal rates of drug and diagnosis yields the relative reporting ratio; pairs of drug and diagnosis with the highest relative reporting ratio are expected to have therapeutic relationships. We also report the performance if the ratios are adjusted using the Gamma-Poisson Shrinker method,[23] an empirical Bayes approach which has been used to mitigate the influence of low counts in these two-by-two tables.

As another measure of the quality of the embeddings, we adapted a score from Choi, et al.[11] to quantify for each pair of a drug and an ICD-9 code indication, what percent of the nearest embedding neighbors of that ICD-9 code share that same drug indication: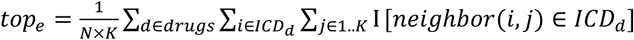

Where:

- *ICD*_*d*_ refers to the set of all ICD-9 codes where drug *d* is indicated
- *neighbor*(*i,j*) refers to ICD-9 code that is the *j*-th nearest embedding neighbor to ICD-9 code *i*
- *K* is a selected number of nearest neighbors to use (we use K = 20)
- *N* is the total number of drug-indication pairs.

A similar measure *frac_e_*, asks: for what percent of the drug-indication pairs, do the K-nearest neighbors include another indication for the same drug: 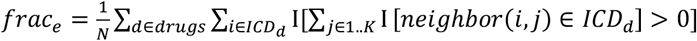

### Creating propensity scores to assess comparability of drugs

Given a set of patients treated with either a drug of interest or a comparator drug, we aim to create a propensity score that models the probability of receiving treatment, as opposed to comparator. This amounts to a large-scale logistic regression to model *p(treated = 1* | *medical history, demographics)*. We used elastic-net regression from scikit-learn to fit the model and estimate the propensity score, with regularization parameter tuning with cross validation. In the logistic regression model, we include the following as predictors:

- Gender
- Age, year, number of prescriptions, diagnoses, and procedures preceding treatment. These are modeled using B-splines with knots placed based on the quantiles of the distribution of these values in the treated population, using the patsy package.
- Indicator variables for presence or absence of each drug, prescription or procedure in a patient’s medical history. Since more recent events would be expected to be more relevant for predicting prescription, we experimented with various features to represent these events. By assessing performance of the resulting classifiers, we settled on representing each event with an indicator variable for if the event is recorded in the previous month before prescription; in the previous year; or ever in patient history

### Implementation and analysis of patient matching

We summarize patients using a weighted average of the embedding vectors corresponding to codes that appear in their history in the time before treatment. Each code is weighted using an exponential decay so that more recent codes count more, and the weights are normalized to sum to 1. This creates a 50-dimensional health summary vector for each patient. This method was chosen because it maximized the accuracy of predicting treatment, but other methods of encoding patient history had similar results. Then we employ a two-step matching scheme that uses Coarsened Exact Matching (CEM) to bin patients, then uses Mahalanobis matching to match patients within bins based on their health summary vectors. For the CEM step we create coarsened patient bins, defined as: gender; calendar year of prescription; age in bins of: 0-5,6-10,11-15,16-25,26-40,41-55,55-70; and number of unique drugs prescribed to that person (divided into percentiles). The CEM has a dual purpose: it ensures we are not creating entirely inappropriate matches (ie, matching children to seniors); and it reduces the number of possible matches for each treated person, making the Mahalanobis matching computationally feasible. Then, we use the sklearn package to perform one-to-one matching of treated patients to comparator patients, based on the Mahalanobis distance between their summary vectors. To compare the Mahalanobis matching against propensity score matching, we use the propensity scores calculated as in the previous section, and perform nearest neighbor matching on the propensity score values, again matching within CEM bins.

Then, we analyze how similar matched pairs are in terms of smoking. There is an ICD-9 code for smoking, but we expect that it only represents some of the information related to smoking status. Therefore, we estimate the probability of smoking by fitting a simple logistic regression model trained to predict whether a person has an ICD-9 code for smoking (code 305.1), given the person’s entire health history, as described in the previous section (removing ICD-9 code 305.1). The top predictors, as expected, include codes for lung diseases and smoking cessation therapies. We use each person’s predicted probability of smoking to evaluate the matching. Note that the same information (including the code for smoking) is provided for training the propensity score function; however this does not guarantee that propensity score matching will match people for probability of smoking. Then, we calculate the correlation between paired patients from the two matching schemes.

## Results

### Overview of indication embeddings

We apply the method outlined in Figure 1 to create a skip-gram set and to learn embeddings. This method yields two types of embeddings: indication embeddings, which signify the health context for a treatment choice; and new-drug (incident drug) embeddings, which represent the treatment choices given in those contexts. The indication embeddings create a unique vector for each possible event in the health history, comprising 3,014 common drugs, 13,090 ICD-9 codes, and 14,187 CPT codes. Figure 2A shows the UMAP[24] placement of all such events, which puts codes with the most similar indication embeddings near each other. Some selected medical events are highlighted. For example, myocardial infarction (MI), medications related to MI, and procedure codes for coronary bypass graft, are all located near each other. Similarly, in a visualization of the new-drug embeddings, drugs from the same REDBOOK therapeutic group appear near each other. This implies that presence of codes for drugs, diagnoses, and procedures all hold information about patient health. In Figure 2B, we group codes into Clinical Classification Software category, for diagnoses, and REDBOOK therapeutic group, for drugs, and then calculate the average indication embedding distance between codes in a pair of groups. For example, at the top, gastrointestinal drugs are near the gastrointestinal system diagnosis codes; at the bottom respiratory systems diseases are near respiratory tract agents.

### Using indication embeddings to predict drug therapeutic uses

Our embeddings were created to summarize the relationship between events in the health history, and treatment choices. A different, but related, task, is prediction of the FDA approved uses of drugs. Similar to Li and Xiao,[6] we use MEDI as our gold standard, as it is based on human curated databases. We examine whether embedding distance between a drug and an ICD-9 code is a good predictor of whether that drug and ICD-9 code have an indication relationship. For comparison, we use the relative reporting ratio: how much more frequently an ICD-9 code is observed before drug prescription, versus the rate expected, and a stabilized version known as the Gamma-Poisson Shrinker. The ROC AUC is .82 for the embeddings, and .80 for both of the disproportionality methods (area under the precision recal curve is .151 and .150 respectively); this is a similar improvement in AUC over disproportionality methods to what Li and Xiao were able to achieve with their method. We also calculate the AUC for predicting the indications for each of 649 drugs and, conversely, for predicting the drugs indicated for each of 1181 diagnosis codes. Grouping these AUC values by REDBOOK therapeutic group, and per CCS class (Figure 3) shows that there is variation in ability of embeddings to predict MEDI relationships. This is expected as some drugs, such as common vaccines, are not strongly tied to any particular diagnosis, and some categories of health conditions, such as congenital anomalies, are not associated with particular drugs.

Comparing the embeddings against previously published embeddings from Choi, et al[11], a study which did not focus on relating indications to treatments, our embeddings are substantially better at representing diagnoses in terms of their relationship to treatments. This is shown in Table 1, where we quantify for pairs of a drug and ICD-9 code indication, what fraction of nearest embedding neighbors of an ICD-9 code share that same drug indication (details in Methods).

**Figure 3.**
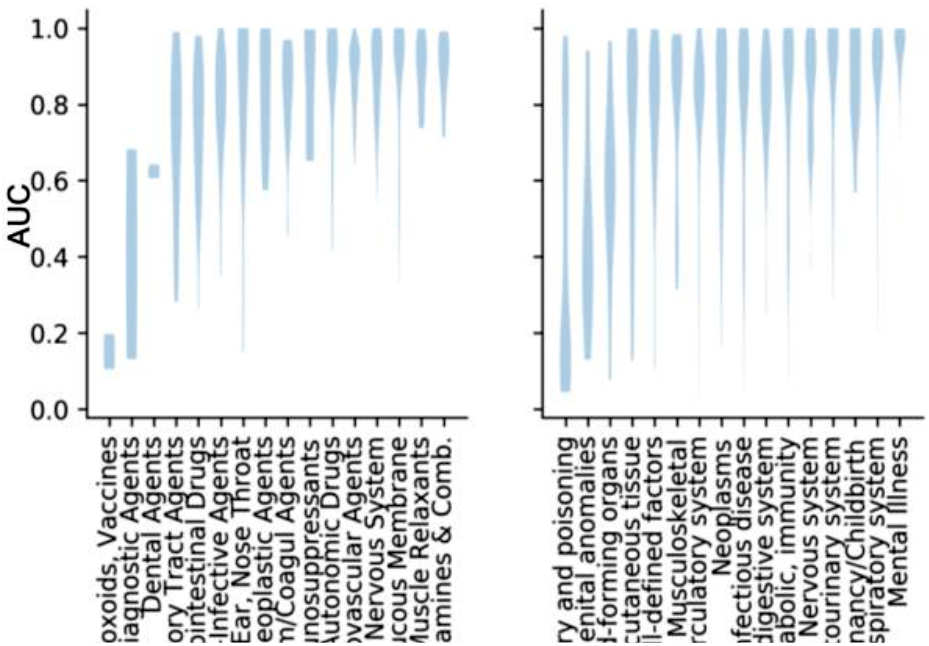
We calculate an area under the ROC for each drug, and for each ICD-9 code. Then we group the AUC results by drug category or ICD code category. Left: Distribution of these ROC values per REDBOOK therapeutic group. Right: per ICD-9 codes in each CCS grouping.

**Table 1:** Comparison of indication embeddings to Choi embeddings.

|  | tope | frace |
| --- | --- | --- |
| **Melamed** | 0.83 | 0.25 |
| **Choi** | 0.70 | 0.19 |

Although the results are promising, more interesting is the ability to detect subtleties of the context in which drugs are prescribed. Supplementary Figure 1 shows the variation in the embedding distance between antidepressants and their most associated diagnoses. To simplify visualization, multiple correlated diagnoses have been collapsed into clusters, and each cluster is represented by the disease nearest its centroid, in Figure 4. Antidepressants include drugs with a range of mechanisms of action, and they are prescribed for diverse reasons. Most have side effects that can influence prescription choice. For instance, tricyclic antidepressants (TCAs) are no longer the standard of care as a first-line treatment for depression (while selective serotonin reuptake inhibitors (SSRI), serotonin norepinephrine reuptake inhibitors (SNRI), mirtazapine, and bupropion are preferred[25]). TCAs can be a second-line depression treatment, but another common use is for chronic pain.[26] This is reflected by the closer embeddings distance of these antidepressants to neuropathy and spine diseases, as well as Irritable Bowel Syndrome (IBS). Pediatric neuropsychiatric diseases are closest to fluvoxamine, while mirtazapine is closest among antidepressants to neurodegenerative diseases of the elderly, which are often associated with depression. Indeed, mirtazapine is a widely used in populations with dementia.[27] This shows how the embeddings represent the complex medical context in which drugs are prescribed.

**Figure 4.**
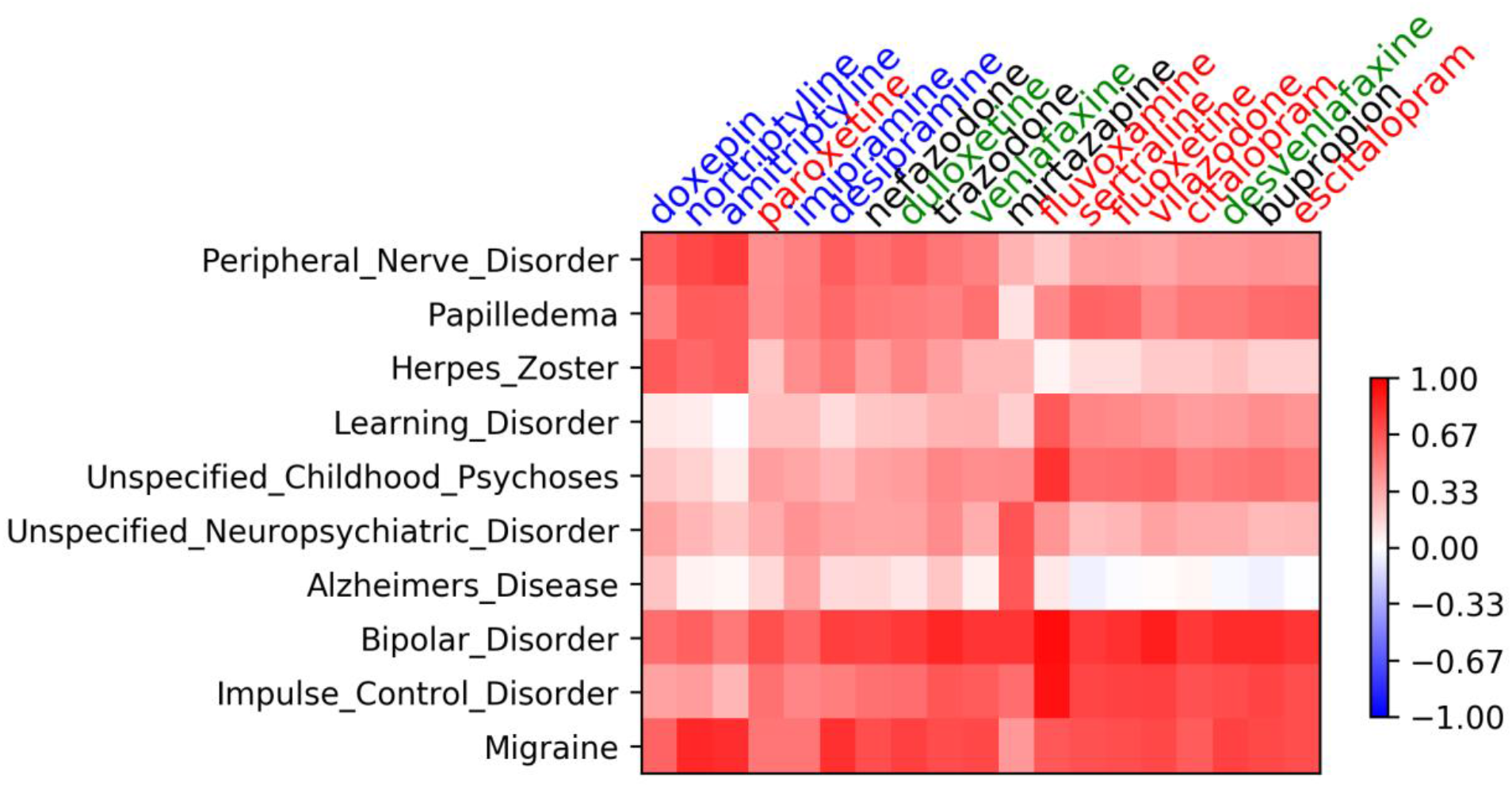
The dot-products between drug vectors (columns) and groups of diagnosis codes (rows) for the diagnoses most related to the antidepressants. Because multiple health conditions are very similar (such as ADHD and Learning Disorders) diagnoses have been clustered and one representative diagnosis for each cluster is shown. Antidepressants are colored by class: TCA are blue; SSRI red; SNRI green; other black.

**Figure 5.**
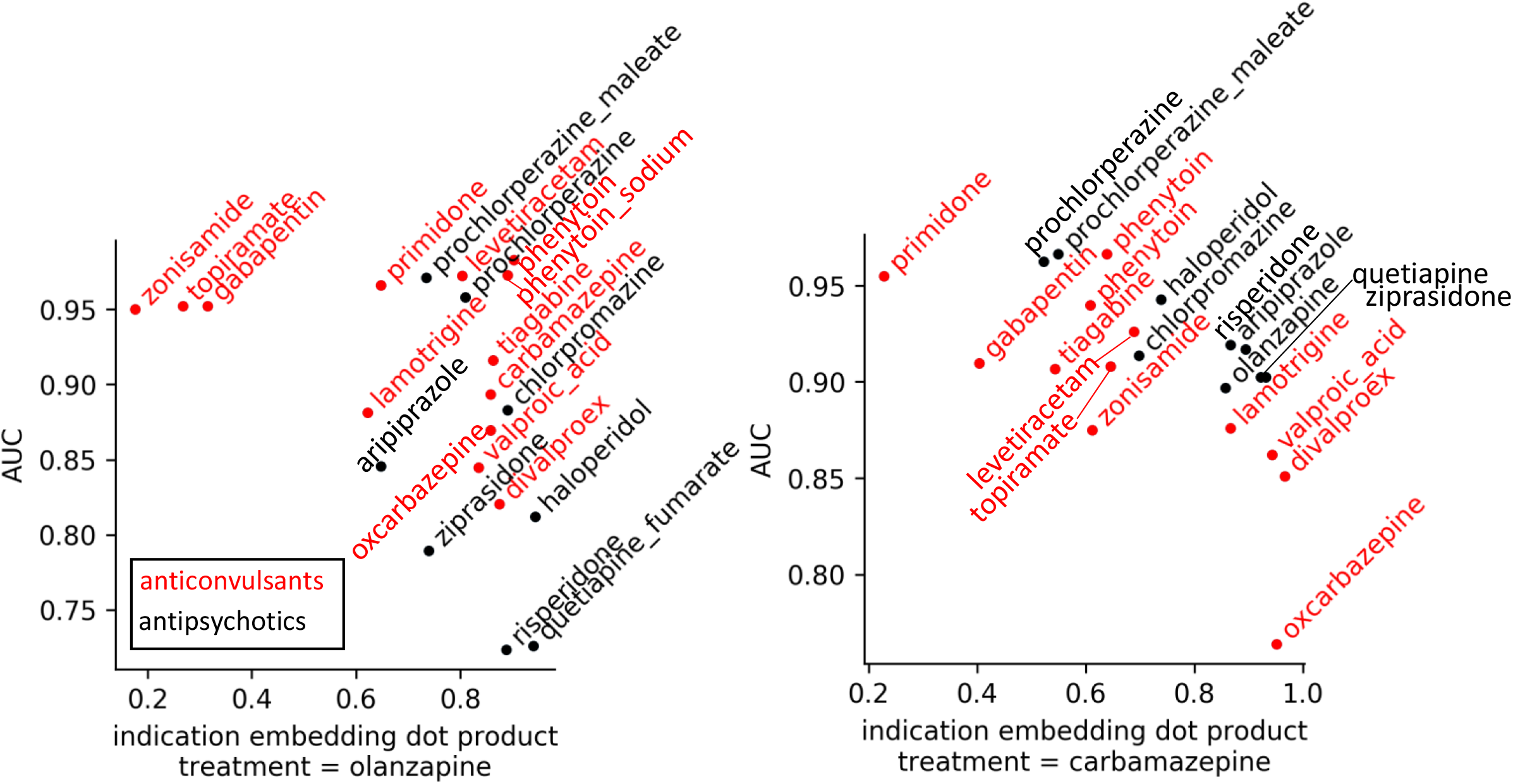
We compare the embedding similarity (x-axes) against the AUC of the propensity scores (y-axes) for finding comparator drugs for olanzapine, left, an antipsychotic (Spearman correlation = −0.58) and carbamazepine, right, an anticonvulsant (Spearman correlation = −0.49. Lower AUC indicates that our propensity model is less successful in separating treatment and comparator drug patients, suggesting the patients are more similar.

### New-drug embeddings point to appropriate comparator drugs

We now turn to the application of the embeddings for drug safety studies, in particular, cohort studies. These study designs typically compare outcomes in the pool of new users of a drug of interest, against the people who took a comparator drug. Selecting an appropriate comparator drug is the first step toward removing differences between the exposed and comparator cohorts that can confound the results of cohort studies. Typically, a medical expert selects the best comparator, but here we assess the performance of embeddings for this task. Drugs that are most comparable should be given in the most similar medical contexts, such as two drugs for the same condition. This is exactly the information that we have shown above is captured in our embeddings. Therefore, we evaluate whether drugs with closer new-drug embeddings are more comparable. We evaluate comparability of a pair of drugs as the comparability between two cohorts of patients that are defined by that pair of drugs. The first of these cohorts comprises patients who received the treatment, and the second cohort contains patients on the comparator drug. The comparability of these cohorts can be assessed using the performance of a classifier to predict whether each person in the cohort study received the treatment or the comparator drug.[28] The easier it is to distinguish these two cohorts, the less comparable they are. Thus we fit the propensity score, p(treated | medical history), a classifier that summarizes the probability of each person getting the treatment of interest. In very comparable cohorts, the propensity to be treated, given a particular medical history, should be similar to propensity to get the comparator drug, which would result in a low area under the ROC curve (AUC) for the propensity score model. Conversely, less comparable cohorts are easily separated and have a high AUC. Figure 5 shows that closer distance between embeddings vectors is associated with lower AUC, which indicates that drugs with closer embeddings indeed are more appropriate comparator drugs for a given treatment. Our embeddings, then, can be used to suggest the most appropriate comparator drug among a number of choices, even in the complex case of neuropsychiatric drugs with multiple indications, and they even have the potential to allow automatic selection of comparators.

### Using embeddings to match patients in a cohort study

Choosing appropriate comparator drugs is only the first step in removing confounding differences between cohorts. This is typically followed by further adjustments, such as propensity score matching or propensity score weighting. Weighting can result in extreme values if very incomparable patients are included. Matching patients again forces researchers to confront the high dimensionality of the health care record. Exact matching of health histories is impossible. One alternative is coarsened exact matching,[22] but that method requires that researchers choose only a few important variables to match on, and it is not easily extended to the high-dimensional setting. In contrast, propensity score matching reduces the high-dimensional health history to a single dimension: the propensity for treatment, which is the probability of treatment given health history. This reduction can result in loss of valuable information regarding patient state, resulting in matching dissimilar people.[29] Mahalanobis matching instead pairs patients based on their distance, given a vector of patient characteristics; but this matching, like coarsened exact matching, does not extend to the setting of sparse uninformatively coded data. Therefore, we experiment with a new matching method. We summarize each patient’s health status as a weighted average of their embedding vectors. Then, we perform a two-step matching strategy that first, matches patients on coarsened versions of age, year, and number of prescriptions, as well as gender, then second, within these matched bins of patients, performs Mahalanobis matching on the lower dimensional *health summary vectors*.

An example of the difficulty of patient matching in observational cohort studies can be found in a comparison of two common antidepressants: bupropion and trazadone. In a study estimating the effect of bupropion on some outcomes, each person taking bupropion could be matched to a person taking trazodone using the propensity score. Each drug has a side effect risks and contraindications that can influence treatment choice and thus propensity for treatment assignment. In addition, bupropion is approved not only for depressive disorders, but also for smoking cessation. Since smoking can cause a number of downstream effects on health, it is desirable to match bupropion-users who are smokers to trazodone users who are smokers. In Figure 6 we show the UMAP visualization of the health summary vectors for a set of people on bupropion versus trazodone, within one coarsened bin. We also include people taking varenicline, a smoking cessation drug, for comparison. Since varenicline users presumably smoke, people nearer to the varenicline population should be more likely to be smokers. Naturally, the Mahalanobis matching on health summary vectors (Figure 6B) results in points from closer parts of the plot being matched. Thus it appears bupropion-takers who are similar to the varenicline population are matched to trazodone-takers who are also similar to the varenicline population (that is, smokers). To see whether this results in smokers being matched other smokers, we create a simple score to summarize probability of smoking in each patient (see Methods) and calculate how similar this score is between matched patient pairs, using Spearman correlation. The correlation is 0.57 for the Mahalanobis matched pairs, and 0.34 for the propensity matched pairs. This shows that Mahalanobis matching on health summary vectors improves matching on specific health risk factors. To demonstrate that this effect is more general, we construct a number of matchings between pairs of antidepressants, and perform the same evaluation to estimate how well matched these patient sets are on a number of different disease conditions (Figure 6C). Matching on embedding vectors consistently results in matched pairs with a more similar health status, across a number of health conditions. Such a matching can create more comparable cohorts, which can then be used with traditional methods in cohort studies to estimate the effects of drugs.[30]

**Figure 6.**
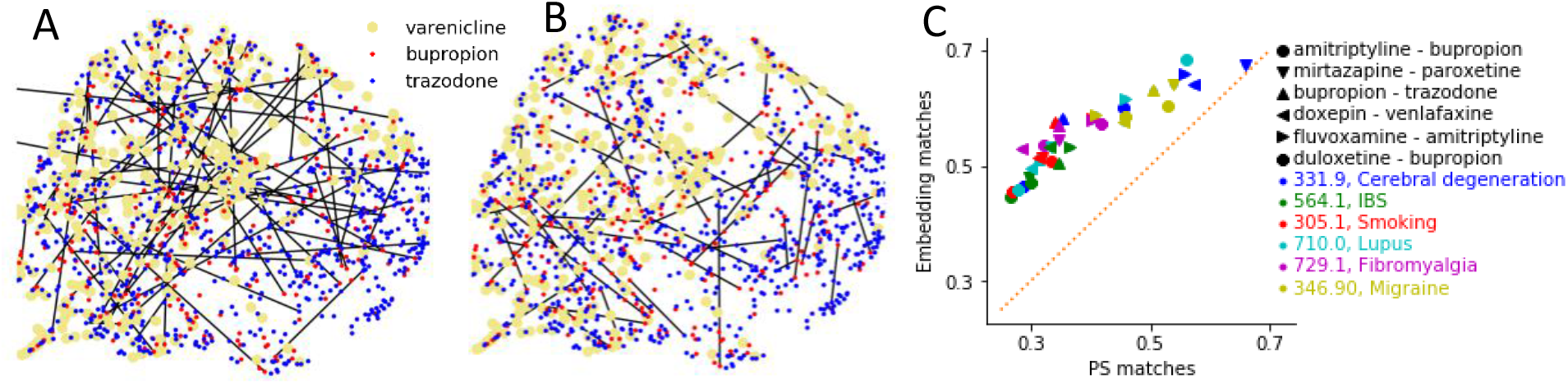
Visualization of the matching of bupropion to trazodone patients. Each point is one person on bupropion, trazodone, or varenicline, using a UMAP visualization of people’s health summary vectors. The location of the yellow points shows that varenicline-takers cluster in the upper left, so people who are on the bottom/right probably are not in need of smoking cessation. The same points are shown in A and B. A. Lines connect 57 pairs matched by propensity score matching. B. Lines connect 61 pairs using Mahalanobis matching on health summary vectors. In the Mahalanobis matching (B) bupropion-takers in the smoking area are more likely to be matched with trazodone-takers in the smoking area. C. We estimate for each patient taking any of nine antidepressants the probability that they have each of six health conditions. Then, we perform a number of matching experiments (different marker shapes). For each set of matched patients, we calculate how correlated matched pairs of patients are in terms of each disease presence measure, under the propensity matching scheme (x-axis) and Manalanobis matching scheme (y-axis). Dashed line indicates identical correlations.

## Discussion and Conclusion

We have shown that indication embeddings form representations of medical events that reflect the complexity of health care. While the indication embeddings work well for predicting known drug indications, their true utility is in their ability to summarize the health status of a patient implied by presence of a given code in a medical history. The decision to prescribe a medication is a complex consideration of personal history, tolerability of side effects, and variation in severity of disease, making such a representation desirable. The motivation for creating these embeddings is to accelerate discovery in drug safety cohort studies. Cohort study design currently heavily relies on clinician knowledge, but since these embeddings are trained on data that results from similar knowledge, they are a way to describe these considerations in a computable form. We show how the embeddings can identify comparable drugs and suggest more comparable matched populations. These results show how these embeddings can complement the cohort study literature, and could be used in combination with other methods to perform drug safety and comparative effectiveness studies.

Our approach has a number of limitations related to the design choices. In our embedding construction, each medical event occurring in a patient’s history is treated as unrelated to other medical events; an alternative design would instead combine patient history in order to predict the drug, given a combination of medical events. We chose our approach specifically to untangle the relationship between each event and treatment state. Other limitations include the drawbacks of claims data, which is not ideal for research purposes: some diseases have better coverage in terms of codes available than others.

These embeddings are available for public reuse at https://figshare.com/projects/Using_indication_embeddings_to_represent_patient_health_for_drug_safety_studies/67532 and have applications beyond the ones described in this paper. Previous embeddings have focused on modelling disease onset but our focus on modeling health state leading to drug prescription makes these results unique. While we have focused our analysis of the embeddings on the relationships between drugs and diseases, these embeddings also relate to symptoms induced by diseases and procedures associated with drugs. Therefore, our embeddings could be used to discover these associations. As Marketscan aggregates data from many health systems, it should represent the standard of care across the USA. Then, the indication embeddings can also be incorporated into analyses on other USA health data sets which have the same standard codes, but smaller patient sizes, as a way to share information learned on a large national data set. In another area of future work, comparing the embeddings trained on medical data from different health systems will allow comparison of medical decisions. Other types of embeddings may be of interest for future work, such as embeddings trained on the time after drug prescription, which may relate to side effects of drugs. In addition, embeddings that incorporate information about drug mechanism, using resources such as UMLS, could be incorporated to integrate molecular understanding of a drug with clinical usage.

## Acknowledgements

This work was funded by NIH 5K01ES028055. Thanks to Andrey Rzhetsky and members of the Rzhetsky lab for helpful discussions.

