## Supplementary for "Using indication embeddings to represent patient health for drug safety studies"

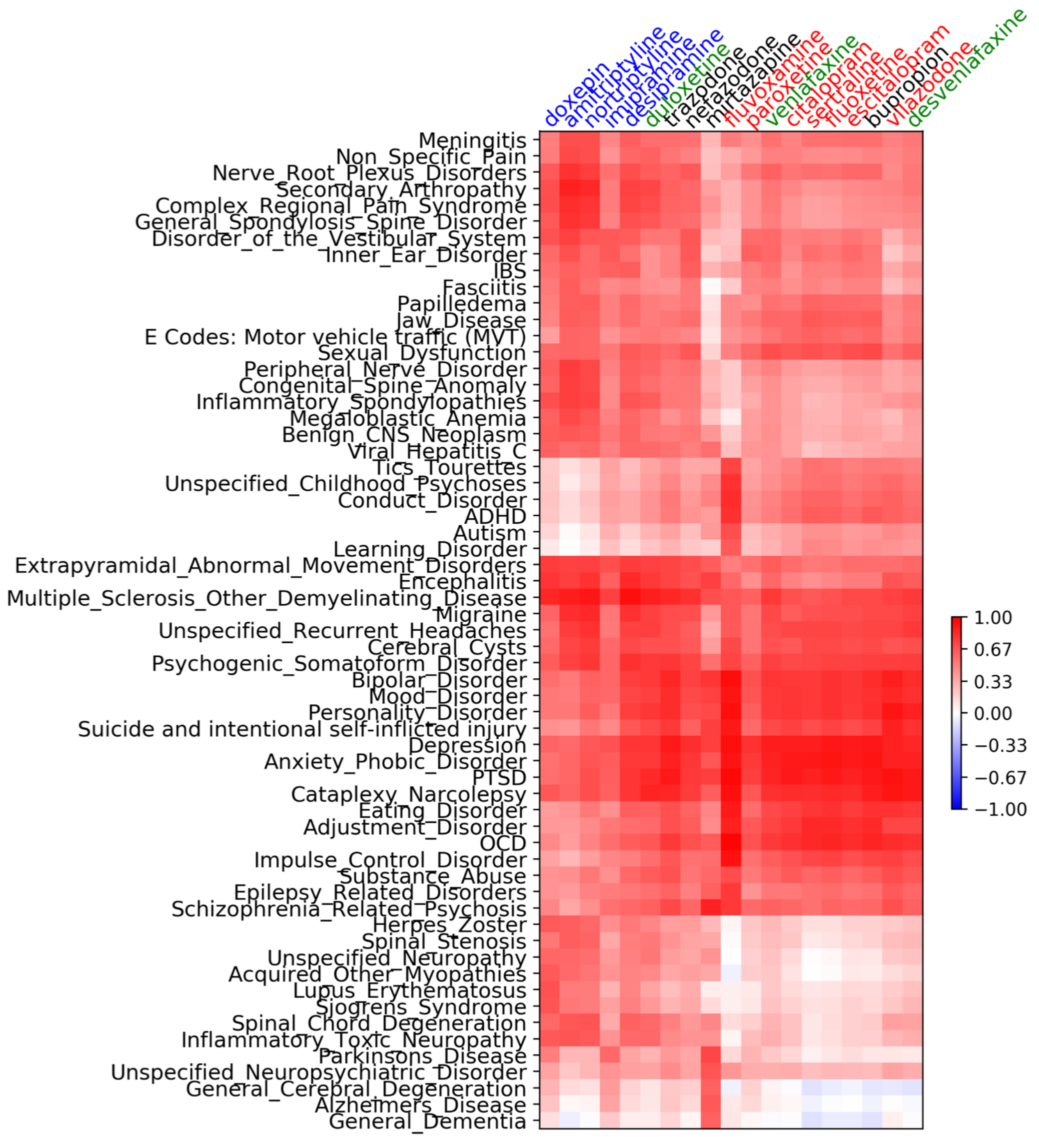


Supplementary Figure 1: The dot-products between drug vectors (row) and groups of diagnosis codes (columns) for all diagnoses most related to the antidepressants. This is an expanded version of Figure 4. Antidepressants are colored by class: TCA are blue; SSRI red; SNRI green; other black

| experiment | chunking | SD window | sampling power | category_cor | auc_medi | top_e | frac_e | correlations_olanzapine | correlations_carbamazepine |
|---|---|---|---|---|---|---|---|---|---|
| **0** | **4** | **40** | **0.5** | 0.98 | 0.83 | 0.27 | 0.83 | -0.61 | -0.49 |
| **1** | **2** | **40** | **0.4** | 0.99 | 0.82 | 0.26 | 0.82 | -0.55 | -0.52 |
| **2** | **1** | **40** | **0.5** | 0.98 | 0.81 | 0.25 | 0.80 | -0.66 | -0.70 |
| **3** | **2** | **20** | **0.6** | 0.93 | 0.82 | 0.28 | 0.82 | -0.84 | -0.39 |
| **4** | **2** | **40** | **0.6** | 0.99 | 0.82 | 0.26 | 0.83 | -0.71 | -0.57 |
| **main text** | **2** | **40** | **0.5** | - | 0.82 | 0.26 | 0.82 | -0.58 | -0.49 |

Supplementary Table 1: Comparing results of 5 different settings of generation of skip-grams (chunking, Standard Deviation window, and parameter used for down-sampling very common codes, as described in the Methods), against the setting used in the main text (bottom row). Each of the embeddings (rows) is compared for: *category_cor*: correlation between the matrix of dot products of category codes in figure 2A in the main text, and the equivalent matrix created for each embedding; *auc_medi,* $top_e$ and $frac_e$ as described in the main text; and the correlation coefficients between drug dot products and propensity score AUCs, as in Figure 5.
